# Improved estimation of SNP heritability using Bayesian multiple-phenotype models

**DOI:** 10.1101/139162

**Authors:** Najla Saad Elhezzani

**Affiliations:** Department of Medical and Molecular Genetics, King’s College London; Department of Statistics, London School of Economics and Political Science; Department of Statistics, King Saud University

**Keywords:** SNP heritability, multiple-phenotype, overfitting, Bayesian analysis

## Abstract

Linear mixed models (LMM) are widely used to estimate narrow sense heritability explained by tagged single-nucleotide polymorphisms (SNPs). However, those estimates are valid only if large sample sizes are used. We propose a Bayesian matrix-variate model that takes into account the genetic correlation among phenotypes and genetic correlation among individuals. The use of multivariate Bayesian methods allows us to circumvent some issues related to small sample sizes, mainly overfitting and boundary estimates. Using gene expression pathways, we demonstrate a significant improvement in SNP-based heritability estimates over univariate and likelihood-based methods, thus explaining why recent progress in eQTL identification has been limited.

## Introduction

For many phenotypes, there is a substantial difference between estimates of narrow sense heritability from family studies and variance explained by discovered single-nucleotide polymorphisms (SNPs) from genome-wide association studies (GWAS) [1,2]. This gap is a key component of the missing heritability problem [3]. Existing genotyping technologies have allowed narrow sense heritability to be estimated from unrelated individuals using all SNPs in the genotyping platform (typically most common with a minor allele frequency >0.05) [4]. However, given that we cannot exclude the possibility of existing rare variants with large effects that have not been detected by genotyping arrays, this SNP-specific heritability is only a lower bound of the true narrow sense heritability. Nevertheless, we do not yet fully understand the gap between SNP heritability and the variance explained by replicated SNPs.

Several hypotheses have been postulated and investigated to explain this problem. Recent attempts suggest that previous estimates are biased and that large sample sizes are required to obtain accurate results [5,6,7]. Naturally, violation of model assumptions can result in biased estimates. For example, using a model that does not capture existing epistatic effects will risk biasing the SNP heritability estimates [6]. Moreover, LMM implicitly assumes that all SNPs have an effect on the phenotype as part of the infinitesimal assumption. Violation of this assumption was thought to be a possible source of bias given the widespread belief that the majority of SNPs are null [8]; however, recent studies found that the effect of this assumption is negligible on SNP heritability estimates [7,9]. Furthermore, in twin studies, the phenotypic variation due to any shared environment might be significant. Therefore, a model that accounts for only a unique environment can inflate heritability estimates.

Biased estimates are not necessarily caused by model assumptions violations; they can also be a result of the assumptions of the estimation procedure itself. For example, the “set to zero” convention, a numerical adjustment used to ensure that variance estimates are positive, will upwardly bias the heritability. Additionally, in many cases, when small sample sizes are used, the variance components are inaccurately estimated taking boundary values. These potential sources of bias are all associated with heritability estimates from variance components models.

The restricted maximum likelihood method (REML) [10] is the mainstream method for estimating variance components. It is implemented in a variety of genome-wide software packages, such as the genome-wide complex trait analysis GCTA [11], efficient mixed-model association EMMA eXpedited [12], FAST LMM [13] and the genome-wide efficient mixed-model association GEMMA [14]. All these methods are equivalent in the sense that they are all based on the same classical univariate LMM. Indeed, some of these methods—e.g., EMMA, FAST LMM and GEMMA—even produce identical p-values in genetic association testing applications [14]. However, these methods differ in their computational complexity, with GEMMA being the most efficient in this regard [14].

REML produces unbiased estimates of the variance components if they are allowed to be negative [15]; otherwise, its estimates are very likely to degenerate in the boundary of the parameter space when small samples are used. When a variance parameter is estimated as zero, this should not imply that it is close to zero. Instead, it commonly indicates a large amount of uncertainty about it [16,17,18]. In multiple-phenotype models, the problem with estimates extends to another class of degeneracy, namely, non-positive definite estimates of the covariance matrices. Such estimates not only are uninterpretable but also can result in underestimated standard errors for the fixed-effect part of an LMM [19]. This feature is misleading in GWAS because an SNP of interest is typically tested by modeling its effect as fixed; therefore, an underestimated standard error will lead to overconfidence about the estimated effect.

Multivariate linear mixed models have recently emerged as a tool to increase statistical power by incorporating correlations among multiple phenotypes. Such models can be fitted using, for example, multi-trait mixed-model MTMM [20] and GCTA [21], both of which are limited to bivariate phenotypes. A popular multivariate method that extended the number of phenotypes to more than two was recently proposed by Zhou and Stephens and implemented in GEMMA software [22]. Both the univariate and multivariate versions of GEMMA are widely used in genetic epidemiology. Therefore, we use them as our benchmark given that they have one advantage over the aforementioned methods: speed [14, 22]. GEMMA relies on the maximum likelihood (ML) method including its restricted version. In this study, however we propose that an improved estimation is obtained using a full Bayes approach, specifically, the use of inverse Wishart (IW) prior for the covariance matrices and a diffuse normal distribution on the covariate coefficients.

Two key problems are addressed by our approach. First, it takes into account the tendency of the ML or REML estimates of the covariance matrices to be non-positive definite even when the number of phenotypes is not very large. Our approach overcomes this problem by adding an extra level of variability to the model through the assignment of a non-informative IW prior that allows the data to dominate while guarding against positive definiteness problems. Second, and as a result, the marginal heritability estimates are more efficient in terms of certainty than those from existing univariate and likelihood-based methods.

The outline of this article is as follows. First, we provide some definitions and notations about the matrix-normal distribution. Second, we discuss the most commonly used model for multiple phenotypes and subsequently state the definition of marginal SNP heritability. Third, we unravel the equivalence between the multivariate model under study and the multivariate ridge regression. This equivalence indicates that the model has the advantage of including all tagged SNPs while accommodating inevitable correlations among them (linkage disequilibrium). The ridge representation is used further to (1) explain the degeneracy problem associated with estimates of the covariance matrices in genome-wide studies and (2) provide a fast evaluation of the posterior distribution of the SNP effect sizes, which can subsequently be used for predictions as model checking. In the last part of the Methods section, we present the complete Bayes model and its simplified form, which facilitates the use of many “off-the-shelf” Bayesian software programs. To show the benefits of the Bayesian multivariate model, we apply it to the expression of genes involved in a breast cancer pathway. We perform two types of comparisons: Bayesian versus frequentist approaches and univariate versus multivariate approaches. Finally, we use a scaled version of the IW to assess for prior sensitivity.

## Methods

### Definitions and notations

The matrix-normal distribution is a generalization of the multivariate normal distribution, which allows us to model correlations among and within subjects [23]. The probability density function for the random matrix X (d×n) that follows the matrix-normal distribution with mean matrix M (d×n) column covariance matrix A (n×n) and row covariance matrix B (d×d) denoted by *X* ∼ MN_*n,d*_(*M, A, B*) has the following form:

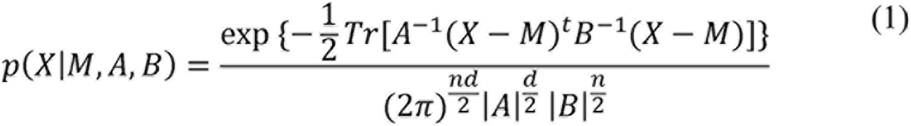

Its expected value and second-order expectations are given by E[X]=M, E[(X – M)(X – M)^*t*^] = B Tr(A) and E[(X – M)^*t*^(X – M) = A Tr(B), respectively.

One way to understand how the matrix normal generalizes the multivariate normal distribution is to assume we have n 1-dimensional variates that are independent and identically distributed as normal with zero mean and variance 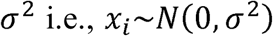. This can be written equivalently as a multivariate normal distribution 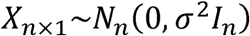. Now, assume we have n d-dimensional variates that are independent and identically distributed as multivariate normal with zero mean and covariance matrix B, i.e., the vectors *X_i_* ~ *N_d_*(0, B). Because these variates are independent, concatenating them will result in a vector with a block diagonal covariance matrix 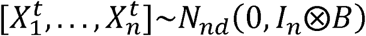, which is itself equivalent to 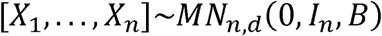.

### Multiple-phenotype model

We consider the matrix-variate model given by

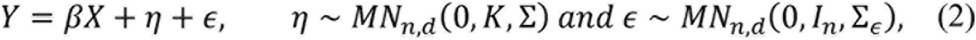

where n and d are the number of individuals and phenotypes, respectively. Here, Y is a d×n phenotypic matrix; X is a k×n matrix of covariates, such as age and sex; and *β* is a d×k matrix of corresponding coefficients. *η* is a d×n matrix of random effects that is independent of the d×n matrix of errors *ε*. The random effect term is used to model any correlation between and within individuals. The n×n relatedness matrix K represents the genetic covariance between individuals and is typically estimated in advance using the genotype data of p SNPs and n individuals. In other words, it is the sample covariance matrix based on the genotype matrix Z (p×n) with rows pre-processed to have zero mean and unit variance, *K* = *Z^t^ Z*/*p*. The d×d matrix Σ represents the genetic covariance matrix within individuals. *Σ*_*ε*_ and *I*_*n*_ specify the environmental covariance matrices within and between individuals, respectively.

Below, we state the SNP heritability definition under this model and discuss problems hindering its estimation.

### Marginal SNP heritability

SNP heritability is defined as the proportion of additive phenotypic variance explained by tagged SNPs. The diagonal elements of the genetic covariance matrix Σ represent the polygenic variances of the d phenotypes. Therefore, the SNP heritability of the *i*^*th*^ phenotype according to the multivariate model is defined as follows:

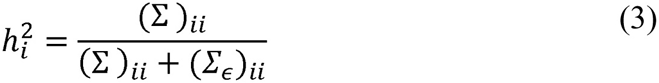

Estimation of Σ and Σ_*ε*_: requires estimation of *d*(*d* + 1) different parameters. Clearly, this number increases rapidly with the number of phenotypes. Such a large number of parameters can make existing algorithms unstable, e.g., by producing covariance matrices that are not positive definite and standard error matrices with large or sometimes uninterpretable entries (NAN). To explain these issues in more detail, it is instructive to first describe the nature of these covariance matrices or, in other words, their relation to SNP effect sizes. To this end, we proceed by writing the matrix-variate model in equation 2 in terms of SNP effect sizes. In statistics, this is referred to as ridge regression.

### Generalized Bayesian interpretation of ridge regression

The Bayesian interpretation of ridge regression assumes that the regression coefficients of a multiple regression are independent and identically normally distributed [24]. Here, we aim to provide a broader Bayesian interpretation of ridge regression in the context of matrix-normal distribution. Consider the matrix-normal regression model of p SNP effects on d phenotypes:

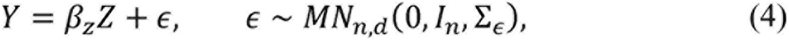

with matrix-normal prior to the effect sizes^1^

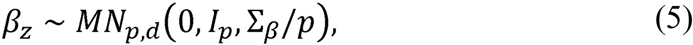

where the 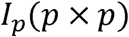 and 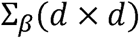 represent the effect size covariances between and within SNPs, respectively. Thus, we are assuming that effect sizes are correlated within SNPs and independent across SNPs. Exploiting the multivariate normal equivalence of matrix-normal distribution, model 4 can be rewritten as follows:

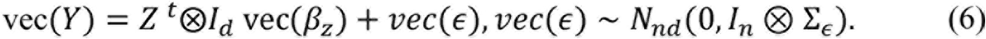

Similarly, the prior on the effect sizes is written as follows:

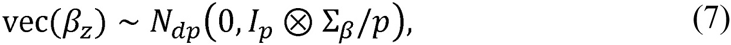

which is itself equivalent to 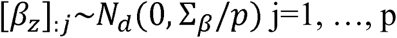.

Here, vec refers to matrix vectorization. Now,

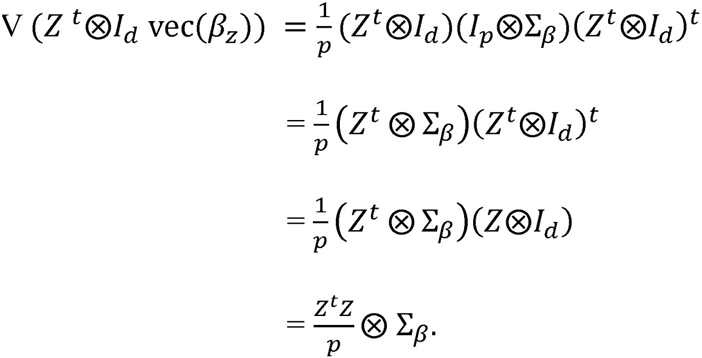

The multivariate normal equivalence of model 2 without βX is given as follows:

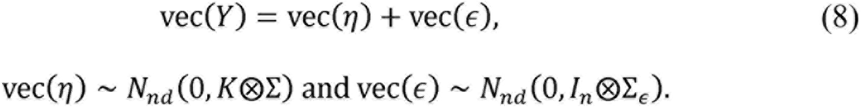

Noting that both vec(*η*) and 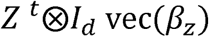 have the same probability model, namely 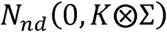, it becomes clear that when the relatedness matrix is estimated using 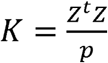, the multivariate ridge regression (equation 4 with 5) is equivalent to the multiple-phenotype model in equation 2. This equivalence shows that the multiple-phenotype model has the advantage of handling linkage disequilibrium in an integrative manner, i.e., without the need for an initial LD pruning step (see [25] for relevant discussion on how ridge regression handles LD).

### Boundary estimation problem

To understand the causes of non-positive definite estimates of the genetic covariance matrix Σ, it is easier to think of model 2 using its equivalent form from the ridge regression representation as follows:

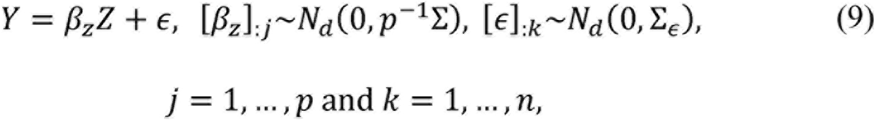

where *β*_*z*_ is the d×p matrix of effect sizes for the d phenotypes and p SNPs. A natural estimator of the genetic covariance matrix would be 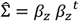. An estimate of this form that is not positive definite may signal a phenotype with zero genetic variance. This occurs when all SNP effect sizes for a particular phenotype are equal. Another common cause is the existence of perfect linear dependency between the SNP effect sizes of different phenotypes. However, given that we know that these conditions are implausible *a priori*, we address the non-positive definiteness problem that corresponds to these factors by taking a Bayesian approach via the assignment of an IW prior to the covariance matrix.

The scaling matrix V of the IW distribution (*IW* (*V*, *v*)) and its degrees of freedom v determine how informative the prior is. For example, the least informative IW is formed by taking the scaling matrix to be the identity matrix and the degrees of freedom to be the least such that the distribution remains proper. Assigning an IW to the covariance matrix is equivalent to assigning an inverse gamma distribution to the variances. To show the effect of the choice of the degrees of freedom on the correlations, 50,000 d-dimensional matrices from *IW* (*I_d_*, *v*) with v=d, d+1 and d+2 were randomly generated (supplementary note)). The least informative IW corresponds to v=d+1, and use of this parameter combination has the effect of setting an approximate uniform distribution on the genetic correlations.

### Simplified full Bayes model

Using univariate LMM, Lippert et al. [13] showed that a spectrally transformed model using a spectral decomposition of the relatedness matrix significantly reduces computational complexity. Similar approaches were subsequently adopted by [26, 22, 14]. Following these developments, we spectrally decompose the relatedness matrix, which allows us to write the matrix-variate model in equation 2 as a multivariate normal model on the transformed data for each individual independently as follows:

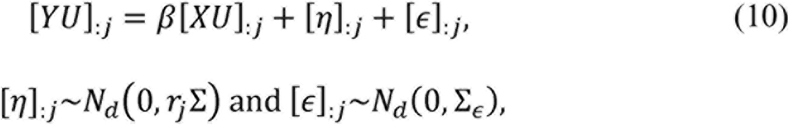

where U is an n×n orthogonal matrix of normalised eigenvectors, and *r*_*j*_ are the corresponding n eigenvalues. Here, [A]_:*j*_ is the *j*^*th*^ column of the matrix A. Eventually, the full Bayes model will have the following hierarchical structure:

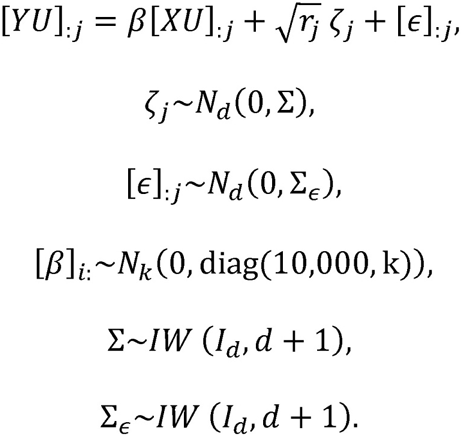

Here, diag (10,000, k) is a k×k diagonal matrix. A BUGS implementation of this model is provided in the supplementary note.

## Results

### The MuTHER Study

One of the main aims of the MuTHER consortium [27] is to quantify the variation in gene expression that is due to genetic factors and ultimately provides insights into the mechanisms underlying the disease susceptibility of associated SNPs. To this end, genome-wide expression profiles (Illumina HT-12v3 Chip) and genome-wide association data (Illumina 610k or 1M chip) were obtained from three tissues adipose, skin and lymphoblastoid cell lines (LCL) from blood samples of 856 Caucasian female twins aged 38.7 to 86.6 years and living throughout the United Kingdom. All recruited females were from the TwinsUK adult registry [28, 29].

We used the SNPs from the original study, which were filtered at a MAF >0.01 and IMPUTE info value >0.6, resulting in a total number of SNPs p=2,238,276. We also used the filtered list of probes given by [27], which excluded polymorphic probes and probes mapping to multiple genes or to genes of uncertain function given that these data are difficult to interpret.

Our model is not tailored for twins in the sense that it does not consider the shared environment, which can result in inflated heritability estimates. Accordingly, we chose to analyze one individual of each twin pair in addition to the available singletons, resulting in a number of individuals suitable for the analysis (N=446). There were no differences in batch effects after removing the twin structure; therefore, only age was included as a non-genotype covariate.

The expressions were downloaded directly from ArrayExpress, and access to the genotypes and covariates was granted from the TwinsUK Steering Committee.

### Heritability Estimation

To illustrate the benefits of our Bayesian multivariate approach, we applied it to a pre-defined gene set from the MuTHER project, namely, genes in a breast cancer pathway [30]. We chose 20 filtered genes comprising d=30 filtered probes. We implemented the simplified full Bayes model as a module for the existing and widely used Bayesian analysis software rjags and coda [31, 32]. We performed two types of comparisons to characterize the variability in the heritability estimates: Bayesian versus frequentist approaches and univariate versus multivariate approaches. In each scenario, confidence and credible intervals of the heritability estimates were inspected to assess their uncertainty.

For univariate likelihood-based analysis, GEMMA software was used to obtain heritability estimates, which also facilitates the use of Wald’s method to compute confidence intervals. The model was separately fitted to each probe when d=1 using its gene expression as a response variable and the genotypes as explanatory variables modeled via a random term. Figure 1 presents the Wald’s confidence intervals for the univariate likelihood-based heritability of each probe. The confidence intervals are wide. In many cases, the intervals spread beyond the parameter space (e.g., including negative values for heritability), which make them difficult to interpret.

**Figure 1.**
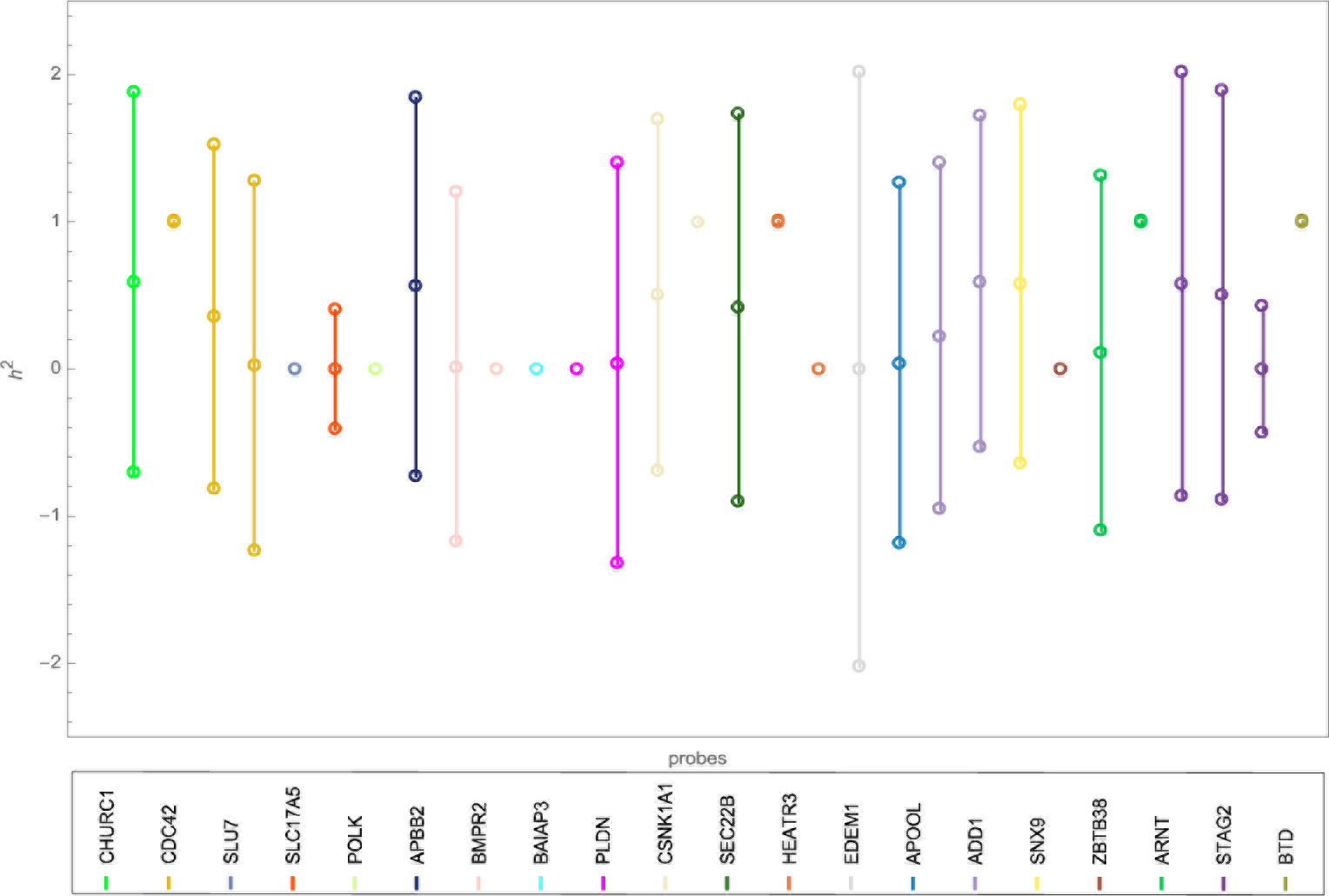
Interval plot of the heritability of 20 BC genes (30 filtered probes) in the LCL tissue using Bayesian univariate analysis with a diffuse gamma prior on the variance components.

For a better characterization of the variability in the heritability estimates, a Bayesian univariate model was used. The model is based on a diffuse gamma prior for the scalar precisions: G(0.001,0.001) with a unity mean and variance of 1000. The Bayesian estimates from the univariate analysis differ significantly from their REML counterparts (figures 1 and 2) as both remain susceptible to variability. Thus, the variance/uncertainty remained large despite the very large number of iterations (see [33] for relevant discussion). Convergence is further discussed below.

**Figure 2.**
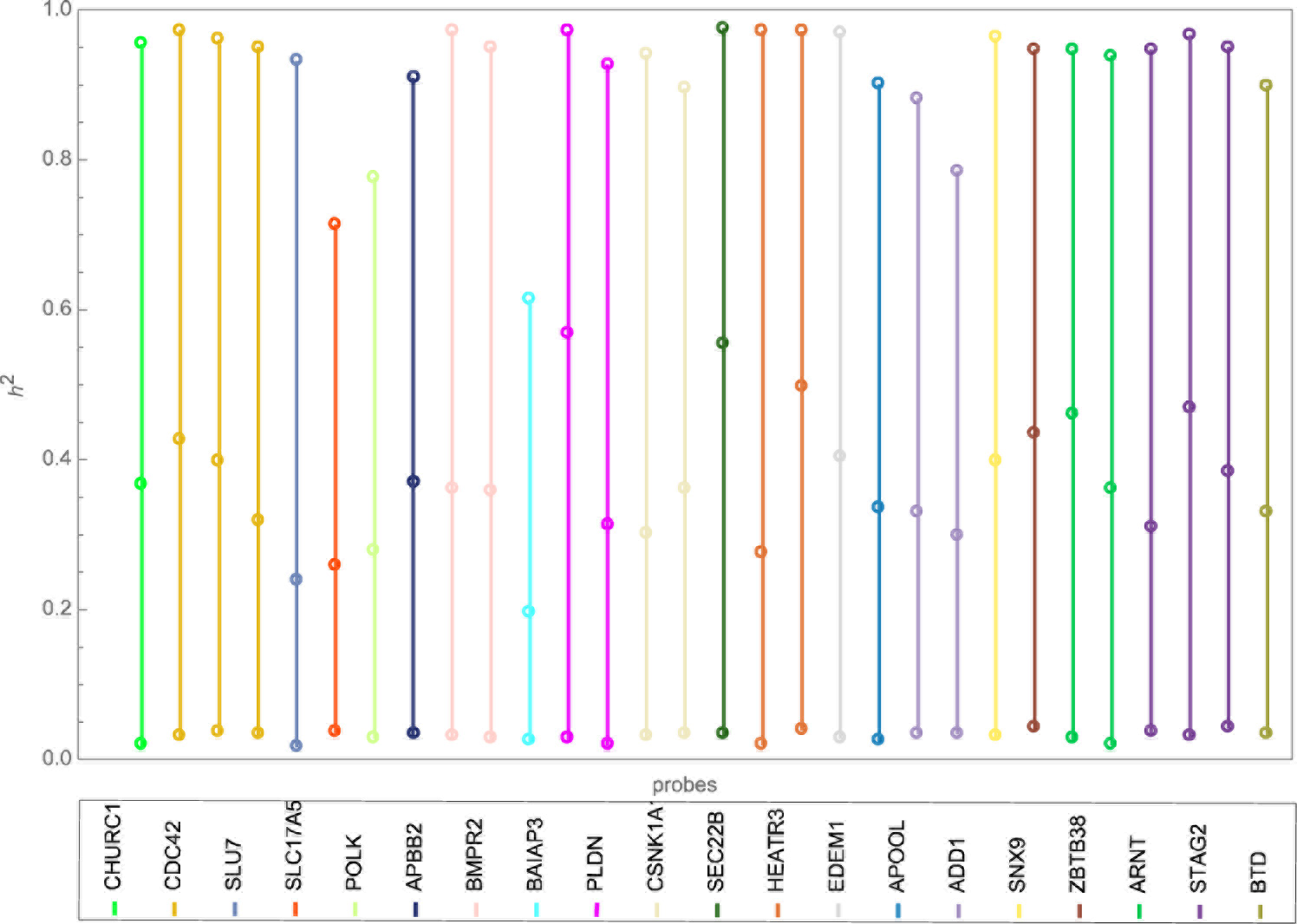
Interval plot of the heritability of 20 BC genes (30 filtered probes) in the LCL tissue using Bayesian univariate analysis with a diffuse gamma prior on the variance components.

Although GEMMA can theoretically be applied to any number of phenotypes, when we attempted to fit 5 probes from the breast cancer pathway using the filtered set of individuals (N=446), the numerical algorithm failed to produce valid standard error matrices. In addition, the covariance matrices were not positive definite because of the small sample size and large covariance matrices. We overcame this problem by assigning the IW prior, as described above.

The credible intervals of the heritability of each probe are much narrower under the Bayesian multivariate model (figure 3) than under its univariate counterpart (figure 2), suggesting very little uncertainty regarding the heritability estimates. Our examination of convergence (supplementary note) showed that convergence is not only satisfactorily achieved under the multivariate analysis but also achieved with a shorter MCMC run than under the univariate model. We argue that this significant enhancement in convergence pertains to the extra amount of information used in the Bayesian multivariate model. The genetic correlation matrix does possess useful information. Although 50% of pairwise correlations lie between 0.05 and 0.25, there are pairs of genes with high genetic correlation reaching -0.5 and 0.8 (supplementary note).

**Figure 3.**
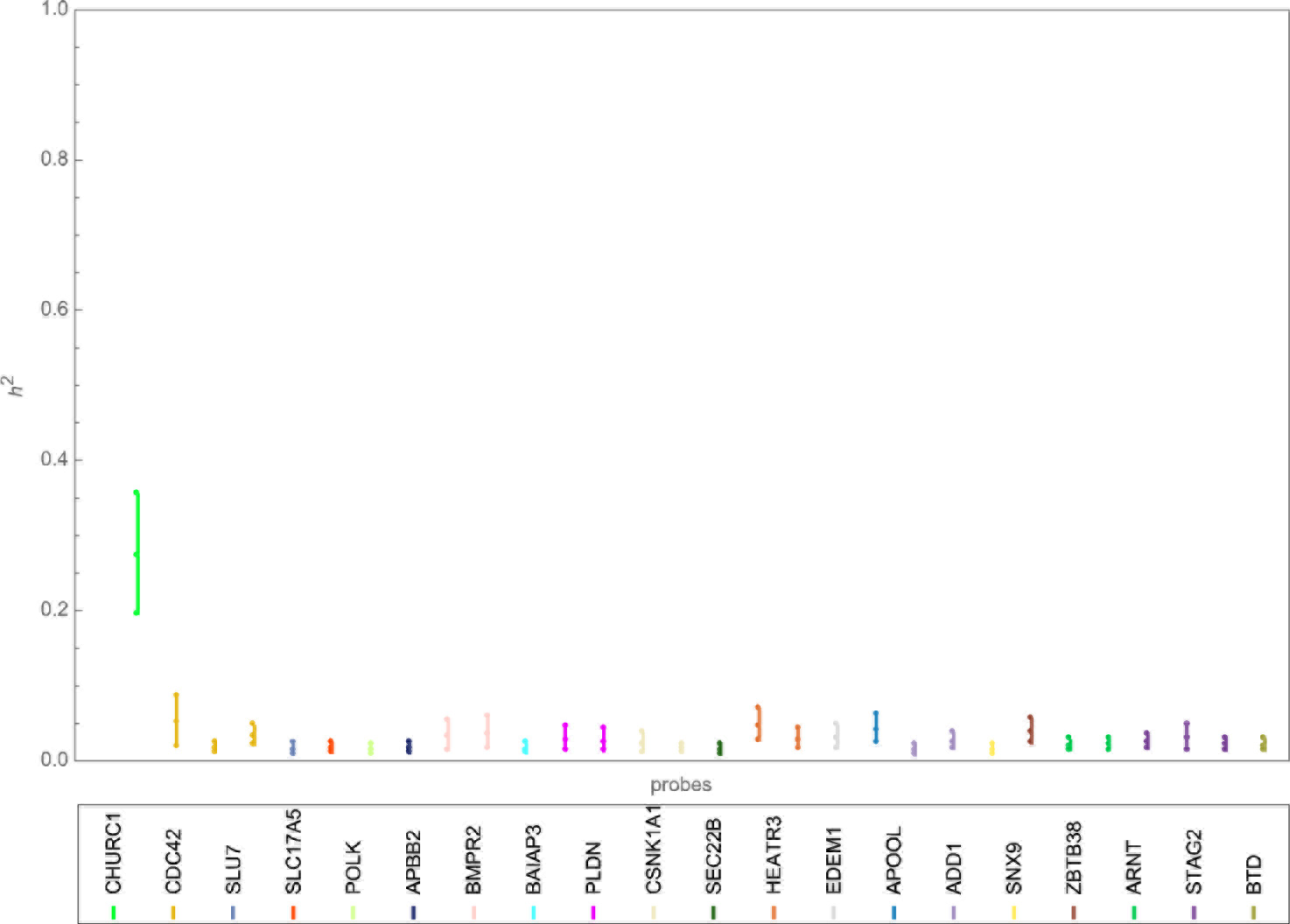
Interval plot of the heritability of 20 BC genes (30 filtered probes) in the LCL tissue using Bayesian multivariate analysis with a non-informative IW prior on the covariance components.

Finally, in contrast to both types of univariate analysis, the multivariate full Bayes approach provided an insight into the SNP relevancy in explaining the variation in expression. Specifically, most tested breast cancer genes have negligible heritability (average ~ 0.03). The exception is CHURC1, which has a relatively high heritability of 0.27 with a credible interval (0.2, 0.36)^2^.

### Model assessment

Based on credible intervals, the results from the BC pathway support the idea that Bayesian multivariate models produce more accurate estimates of SNP heritability than classical multivariate and univariate models. It is also clear that the genetic architectures of complex phenotypes are miscellaneous, and no single method will be the most efficient in capturing all of them. The “gold standard” for model assessment and comparison is therefore an improved phenotype prediction using a new data set. This method can be performed first by estimating the effect sizes using the computationally efficient formula for the mode of their posterior distribution

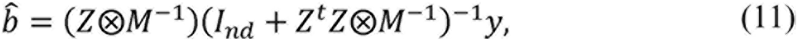

then incorporating the formula to obtain the predicted phenotypic values of the new sample based on its genotypes *Z*_*new*_:

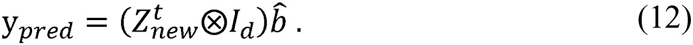

Here, y=vec(Y) and the effect sizes *b* = vec(*β_z_*) are the values estimated from the original data. The prediction algorithm is delineated in the supplementary note.

In our example, phenotype prediction as a model checking technique has its pitfalls. In the breast cancer pathway, our method produced expression heritability estimates that are close to zero. The exception was CHURC1, with a relatively higher heritability. Therefore, any attempt to predict the expression using SNPs is expected to fail because the heritability (variance explained by genotyped SNPs) is very low. Accordingly, we had to resort to alternative approaches in an attempt to provide evidence in support of our SNP heritability estimates. The approaches were (1) finding literature that could support or refute the results and (2) using more flexible prior specifications.

### Literature support

To trace back the heritability estimates, we looked at previously reported *cis* eQTLs for the 20 genes from the MuTHER study tested here. According to Grundberg et al. [27], there are 196 SNPs associated with CHURC1 expression, i.e., with p-values < 10^−8^; however, the other 19 had no reported eQTL, supporting the finding that their expression has limited SNP heritability. To determine whether the same conclusion can be drawn using a different analysis, we used univariate GEMMA to scan chromosome 14 for association with CHURC1 expression and identified 180 SNPs with p-values < 10^−8^. Given that GEMMA fits an LMM with the fixed-effect part being the genotypes of the tested SNP, proximal contamination [34]; that is the situation when the tested SNP is assumed both fixed and random, can incur power loss. To eliminate this issue, we followed the GCTA approach and excluded chromosome 14 from the computations of the relatedness matrix. In addition to the 180 SNPs identified before the exclusion, only one additional significant SNP was detected because of a slight decrease in most of the p-values after the exclusion. Overall, there is a significant overlap between the SNPs from our analysis and those obtained by Grundberg et al. [27].

It would be interesting to determine what effect excluding the detected eQTLs would have on the heritability estimates. To this end, we repeated the analysis, excluding the SNPs previously associated with CHURC1 expression from the model, and the trace plots for its heritability appeared unstable using the same burn-in period and thinning interval that was used before the exclusion. We therefore explored additional convergence diagnostics, leading to the choice of a million iterations with a thinning interval of 1000 after a similar burn-in period of 150,000. Nevertheless, convergence remained an issue for the heritability of CHURC1. However, the heritability of each of the remaining genes remained close to zero with acceptable convergence.

The lack of convergence for the heritability of CHURC1 after excluding its eQTLs recapitulates the boundary estimation problem: the expected polygenic variance is zero (assuming neither epistatic nor very small effects). However, given that we are sampling from a family of positive definite matrices, the resulting estimate will not be zero, which probably caused the lack of convergence. See the supplementary note for convergence diagnostics.

### Prior sensitivity analysis

Possible arguments against the use of the IW are (1) the uncertainty for all variance parameters are controlled by a single degree of freedom, namely v=d+1, and (2) the lack of independence between the variances and the correlations. To overcome the first issue and alleviate the second, we used a scaled version of the IW.

The scaled IW was first introduced by O’Malley and Zaslavsky [35], and its relative advantage over the unscaled IW was discussed by Gelman and Hill [36]. The basic idea is to decompose the covariance matrix into a diagonal matrix 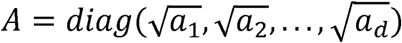 and a matrix V distributed as *W* (*I_d_*, *d* + 1); Σ = *A V A*. This idea implies that Σ~*IW* (*AI_d_A*, *d* + 1), *AI_d_A* = *diag*(*a*_1_,…, *a_d_*). This prior will not shrink the correlations, so their marginal prior will remain uniform when v= d+1. However, the variances now can be estimated more freely from the data. To determine the scaling needed, we added another level of variability by assigning a uniform prior U(0, 100) for each scaling parameter (see supplementary for the relevant BUGS code). The SNP heritability estimates using the SIW are very similar to those obtained using the unscaled method (figure 4), implying that the above concerns that plague the IW do not affect the posterior distributions of the heritability.

**Figure 4.**
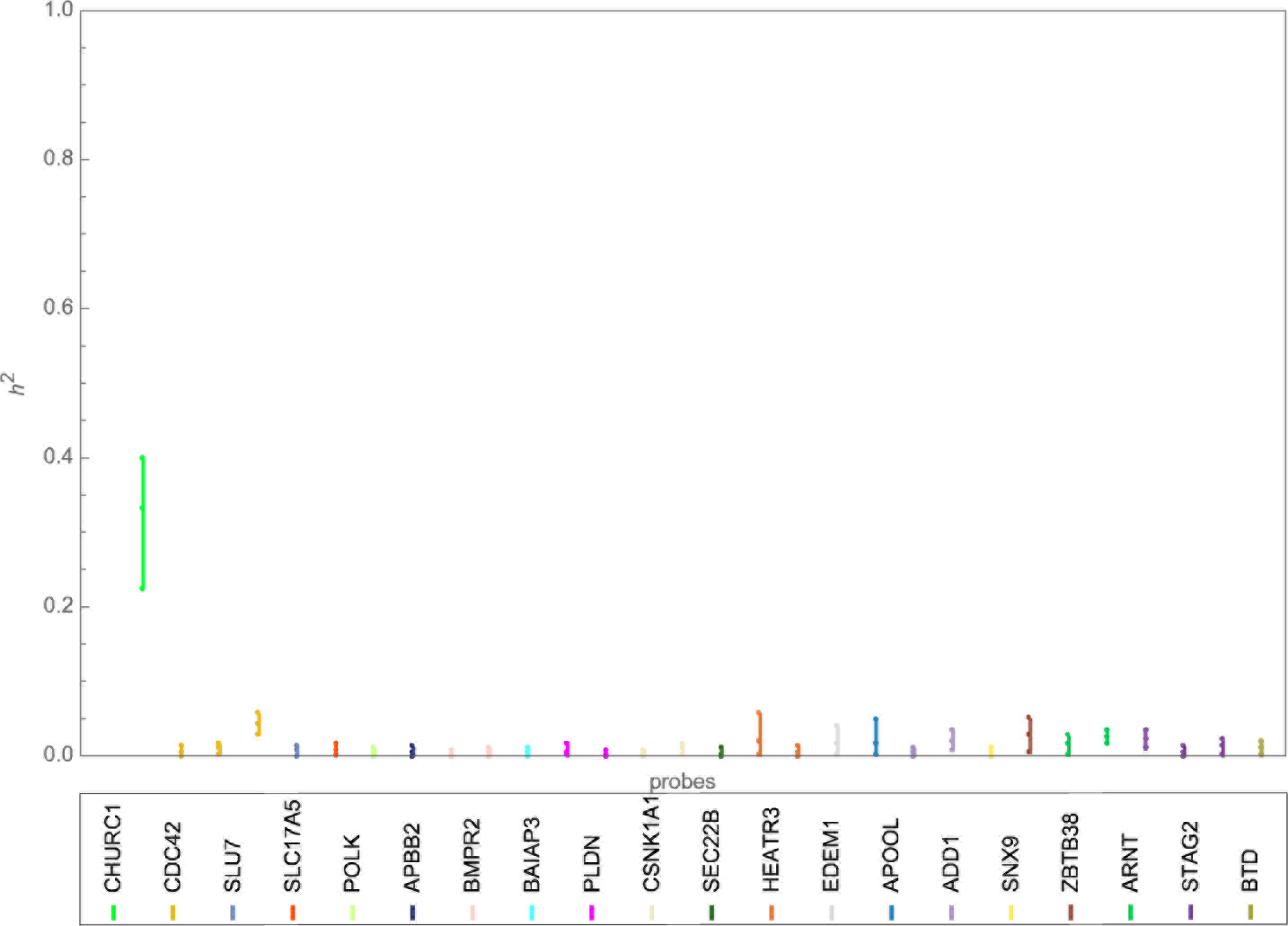
Interval plot of the heritability of 20 BC genes (30 filtered probes) in the LCL tissue using Bayesian multivariate analysis with a non-informative SIW prior on the covariance components and a uniform prior on its scale parameters.

## Discussion

In conclusion, we developed a multivariate Bayesian approach for SNP heritability estimation. Our approach can be seen as an extension of existing multiple-phenotype models, adding an extra level of variability to ensure positive definite estimates of the covariance matrices and ultimately improving estimates of the SNP heritability that can determine how fruitful eQTL mapping strategies can be. From a computational perspective, the simplified form allowed the posterior distribution to be determined at a feasible computational cost using rjags. This feature is advantageous because it saves users from having to write their own MCMC code. However, it should be noted that although rjags performs efficiently for SNP heritability estimation, alternative software might be needed for genome-wide association testing because rjags will be intrinsically slow owing to the number of iterations required by MCMC for the chain to converge. In addition, the multivariate model used herein is based on the infinitesimal assumption; however, the genetic architectures of complex phenotypes are unknown. Therefore, a more flexible prior that fits a wide range of settings may be desirable. A mixture of matrix-variate normal distributions for the effect size matrix is likely to provide a gain in the estimation accuracy and ultimately in the phenotype prediction, representing a promising avenue for future research.

Finally, the results from the SNP heritability estimation of the expression of the tested BC pathway genes, specifically CHURC1, and its eQTLs are provocative and underscore the need to investigate CHURC1 effect on breast cancer status. We have made the first promising steps toward designing a method orthogonal to the one in this paper to investigate the extent to which significant heritability estimates of the expression of BC-related genes will translate into improved predictive accuracy of BC status.

## ACKNOWLEDGEMENTS

The work described in this paper was funded by the Saudi Government as part of the author’s PhD scholarship. The TwinsUK is funded by the Wellcome Trust, Medical Research Council, European Union, the National Institute for Health Research (NIHR)-funded BioResource, Clinical Research Facility and Biomedical Research Centre based at Guy’s and St Thomas’ NHS Foundation Trust in partnership with King’s College London. Many thanks to Wicher Bergsma, Nick Dand and Doug Speed for their useful comments.

## Footnotes

1 Note that *β* and *β*_*z*_ are different. The first value corresponds to the effect sizes of any covariates ozther than SNP genotypes—e.g., sex and age—whereas the second value is specifically for the SNP genotypes, which are stored in Z.

2 These results were recorded after a 150,000 burn-in period using a sample of 5000 resulting from 25,000 iterations with a thinning interval of 5.

